# Integrated Analysis of Tissue-specific Gene Expression in Diabetes by Tensor Decomposition Can Identify Possible Associated Diseases

**DOI:** 10.1101/2022.05.08.491060

**Authors:** Y-H. Taguchi, Turki Turki

## Abstract

In the field of gene expression analysis, methods of integrating multiple gene expression profiles are still being developed and the existing methods have scope for improvement. The previously proposed tensor decomposition-based unsupervised feature extraction method was improved by introducing standard deviation optimization. The improved method was applied to perform an integrated analysis of three tissue-specific gene expression profiles (namely, adipose, muscle, and liver) for diabetes mellitus, and the results showed that it can detect diseases that are associated with diabetes (e.g., neurodegenerative diseases) but that cannot be predicted by individual tissue expression analyses using state-of-the-art methods. Although the selected genes differed from those identified by the individual tissue analyses, the selected genes are known to be expressed in all three tissues. Thus, compared with individual tissue analyses, an integrated analysis can provide more in-depth data and identify additional factors, namely, the association with other diseases.

## 1. Introduction

Gene expression analysis is an important step for investigating diseases and identifying genes that can be used as therapeutic targets or biomarkers or genes that are causes of disease. Although the development of high throughput sequencing technology (HST) has led to continuous increases in the amount of gene expression profile data, methods of integrating multiple gene expression profiles are still being developed. Tensor decomposition (TD) is a promising candidate method for integrating multiple gene expression profiles. Using this method, gene expression profiles from multiple tissues of individuals can be stored as a tensor *x*_*ijk*_ ∈ ℝ^*N*×*M*×*K*^, which represents the gene expression of the *i*th gene in the *j*th individual of the *k*th tissue. TD provides a method of decomposing a tensor into a series expansion of the product of singular value vectors, each of which represents a gene assigned to a specific individual or tissue. For example, by applying the higher-order singular value decomposition (HOSVD) method to *x*_*ijk*_, we can obtain the following:

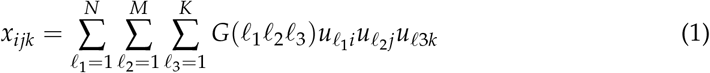

where *G* ∈ ℝ^*N*×*M*×*K*^ is a core tensor, 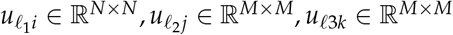 are singular value matrices and orthogonal matrices. We previously proposed a TD-based unsupervised feature extraction (FE) method [1] and applied it to a wide range of genomic sciences. Recently, this method was improved by the introduction of standard deviation (SD) optimization and applied to gene expression [2], DNA methylation [3], and histone modification analyses [4]. Nevertheless, because the updated method was only previously applied to gene expression measured by HST, whether it is also applicable to gene expression profiles retrieved by microarray technology remains to be clarified. In this paper, an integrated analysis was performed by applying the recently proposed TD-based unsupervised FE method with SD optimization to microarray-measured gene expression data for diabetes mellitus from multiple tissues. We found that applying the TD-based unsupervised FE with SD optimization to gene expression profiles from individual tissues can identify diseases associated with diabetes that cannot be identified by the other state-of-the-art methods.

There are multiple benefits in using TD to identify DEGs. First, since it is not a supervised method, it can select DEGs that are biologically more plausible than those selected using supervised methods. This can be explained using the following example wherein the aim is to identify DEGs that are distinct between two classes, e.g., patients and healthy controls. Supervised methods attempt to identify DEGs associated with a smaller divergence within individual classes, whereas TD allows to select DEGs with within-class divergence to some extent (since TD tries to identify the representative state of distinction between two classes). If the representative state is associated with within-class divergence that has biological origins, e.g., age and sex, this divergence should not be penalized. However, supervised methods often do so whereas the unsupervised method allows biological within-class divergence. Second, TD can select more stable DEGs; i.e., those independent of specific sets of samples considered in the analysis. This is because TD attempts to identify DEGs coincident with those of the representative state, which should be robust. Since sub-sampling does not change the representative state drastically, the gene set selected by TD is not altered drastically either. Third, TD can deal with multiple conditions. For example, if gene expression is measured in various tissues of several people, it is natural to format them as gene × person × tissue, which results in a tensor form. We have listed only a few important advantages here. Readers interested in acquiring information on other advantages of TD can refer to our recent book [1].

## 2. Materials and Methods

### 2.1 Gene expression

Gene expression profiles (GSE13268, GSE13269, and GSE13270 [5]) were retrieved from the Gene Expression Omnibus (GEO), and they were obtained from a study of the progression of diabetes biomarker diseases in the rat liver, gastrocnemius muscle, and adipose tissue. Each of these profiles is composed of gene expression profiles from five individuals seen in two strains, Goto-Kakizaki and WistarKyoto, and they include data for three tissues (adipose, muscle, and liver) obtained at five time points after treatment. Three files named GSE13268_series_matrix.txt.gz, GSE13269_series_matrix.txt.gz, and GSE13270_series_matrix.txt.gz were downloaded from the Supplementary Files in GEO.

Gene expression profiles were formatted as a tensor, with *x*_*ijkmst*_ ∈ ℝ^31099×5×5×2×2×3^, representing the expression of the *i*th probe in the *t*th tissue (*t* = 1: adipose, *t* = 2:muscle, *t* = 3:liver) at the *j*th time point for the *k*th replicate and *m*th treatment at *s*th strain. These values are normalized as follows:

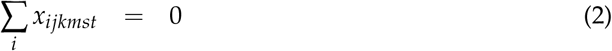

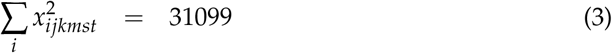

### 2.2 Methods

Figure 1 shows the analysis pipeline. Methodological details can be found in supplementary Information.

**Figure 1.**
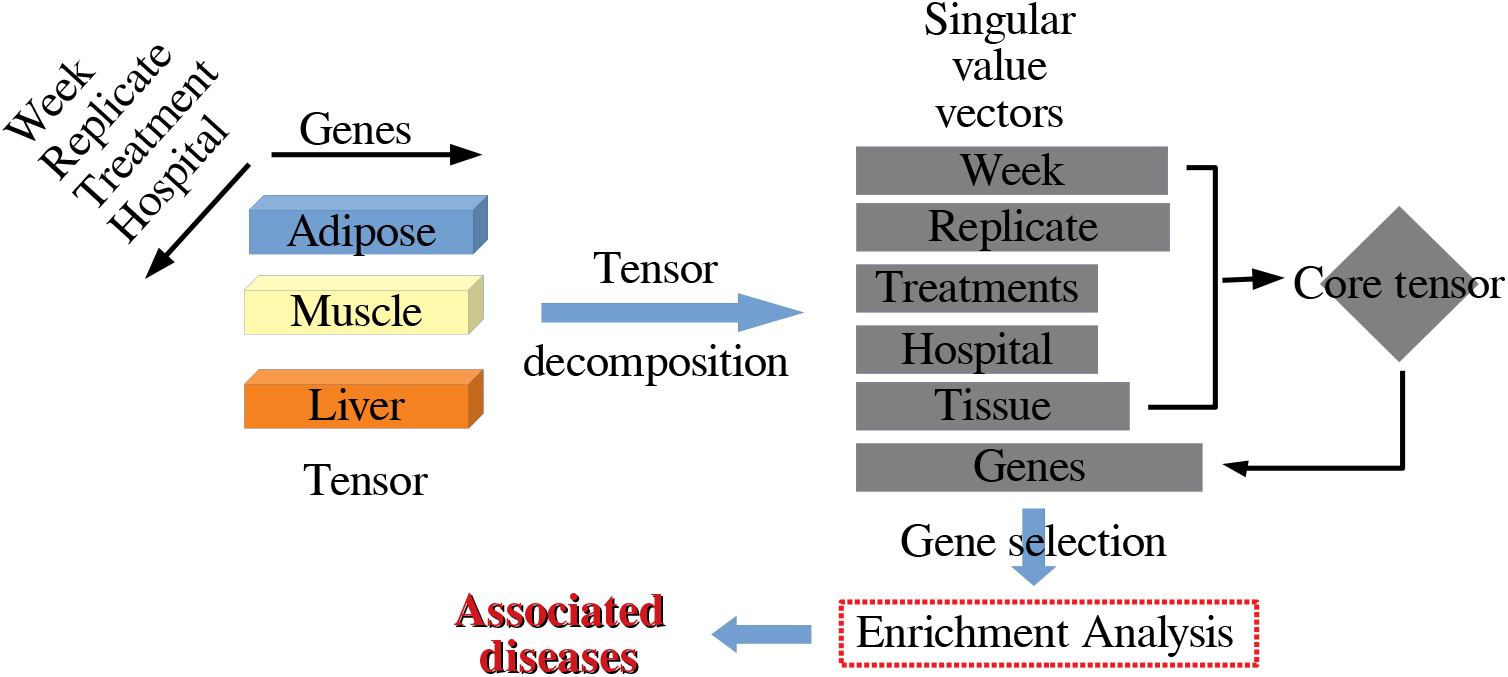
Overall flowchart of the analysis pipeline.

## 3. Results

To validate the selected genes, 2,281 gene symbols are uploaded to Enrichr [6] (For the full list of selected probes, genes and enrichment analyses, check the supplementary materials). Table 1 shows the results of the “KEGG 2021 Human” category in Enrichr. Since none of the terms are related to diabetes except for the top term, i.e., “diabetic cardiomyopathy”, the process initially appears to be a failure. Nevertheless, a number of the identified diseases are deeply related to diabetes mellitus. For example, many neurodegenerative diseases are listed, and diabetes mellitus is widely known to be a risk factor for neurodegenerative diseases [7–11]. Moreover, diabetes mellitus is known to be associated with thermogenesis [12], oxidative phosphorylation [13], and the PPAR signaling pathway [14]. Thus, the proposed method is successful in contrast to the first impression and can identify many diseases associated with diabetes mellitus.

**Table 1.**
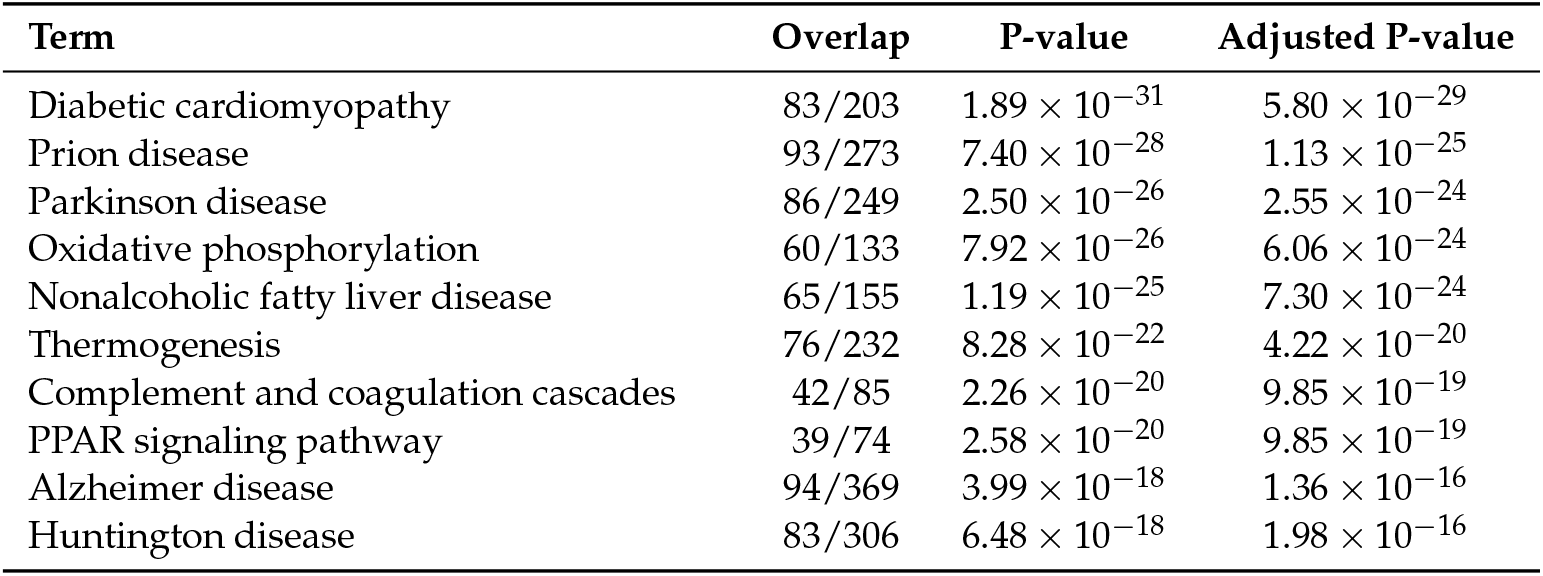
Top 10 “KEGG 2021 Human” category terms in Enrichr.

Table 2 shows the top 10 terms in the category “ARCHS4 tissues” in Enrichr. Remarkably, gene expression is measured for three of the top four tissues. Similar results are found for the “Mouse Gene Atlas” category in Enrichr (Table 3). In conclusion, the proposed method is successful.

**Table 2.** Top 10 terms in the “ARCHS4 Tissues” category in Enrichr.

| Term | Overlap | P-value | Adjusted P-value |
| --- | --- | --- | --- |
| LIVER (BULK TISSUE) | 481/2316 | $3.49 \times 10^{-63}$ | $3.77 \times 10^{-61}$ |
| VENTRICLE | 449/2316 | $1.67 \times 10^{-49}$ | $9.04 \times 10^{-48}$ |
| SKELETAL MUSCLE (BULK TISSUE) | 428/2316 | $2.34 \times 10^{-41}$ | $8.42 \times 10^{-40}$ |
| ADIPOSE (BULK TISSUE) | 410/2316 | $6.46 \times 10^{-35}$ | $1.75 \times 10^{-33}$ |
| MYOBLAST | 409/2316 | $1.42 \times 10^{-34}$ | $3.08 \times 10^{-33}$ |
| SUBCUTANEOUS ADIPOSE TISSUE | 401/2316 | $6.92 \times 10^{-32}$ | $1.25 \times 10^{-30}$ |
| ATRIUM | 366/2316 | $2.38 \times 10^{-21}$ | $3.67 \times 10^{-20}$ |
| HEART (BULK TISSUE) | 363/2316 | $1.53 \times 10^{-20}$ | $2.07 \times 10^{-19}$ |
| HEPATOCYTE | 362/2316 | $2.82 \times 10^{-20}$ | $3.39 \times 10^{-19}$ |
| OMENTUM | 350/2316 | $3.25 \times 10^{-17}$ | $3.51 \times 10^{-16}$ |

**Table 3.** Top 10 terms in the “Mouse Gene Atlas” category in Enrichr.

| Term | Overlap | P-value | Adjusted P-value |
| --- | --- | --- | --- |
| mammary gland non-lactating | 116/201 | $7.92 \times 10^{-64}$ | $7.61 \times 10^{-62}$ |
| skeletal muscle | 229/710 | $5.23 \times 10^{-63}$ | $2.51 \times 10^{-61}$ |
| liver | 243/928 | $3.58 \times 10^{-48}$ | $1.14 \times 10^{-46}$ |
| adipose brown | 148/456 | $5.78 \times 10^{-41}$ | $1.39 \times 10^{-39}$ |
| heart | 154/568 | $2.53 \times 10^{-32}$ | $4.86 \times 10^{-31}$ |
| kidney | 80/554 | $3.98 \times 10^{-4}$ | $5.90 \times 10^{-3}$ |
| osteoblast day 21 | 44/264 | $4.30 \times 10^{-4}$ | $5.90 \times 10^{-3}$ |
| bladder | 33/195 | $1.63 \times 10^{-3}$ | $1.96 \times 10^{-2}$ |
| adipose white | 33/199 | $2.29 \times 10^{-3}$ | $2.44 \times 10^{-2}$ |
| MEF | 45/300 | $3.33 \times 10^{-3}$ | $3.20 \times 10^{-2}$ |

## 4. Discussion

Although the proposed method successfully integrated gene expression data measured in three tissues and identified diseases associated with diabetes mellitus, the identified genes also included genes expressed in all three tissues. If other methods that do not require an integrated analysis can perform similarly, then complicated methods, such as the proposed method, will not be required. To determine whether methods without integration can achieve similar performance, we tested three methods: *t* test, SAM [15] and limma [16]. Since the *t* test and SAM methods cannot simultaneously consider the distinction between the control and treatment as well as the dependent on time, we attempted to identify genes that presented expression differences between the control and treatment (no consideration of time dependence). For more details on how to perform these three methods, check the sample R source code in supplementary materials.

Table 4 shows the number of probes selected by the other methods. These methods select fewer probes than the proposed method (2,542 probes), and the number selected in muscle is relatively low. According to the limma method, only two probes could be selected for muscle; thus, the method was not successful. The integrated analysis likely helped identify more probes, which resulted in more significant enrichment.

**Table 4.** Number of probes selected by other methods.

| Tissue | *t* test | sam | limma |
| --- | --- | --- | --- |
| Adipose | 556 | 773 | 116 |
| Muscle | 100 | 119 | 2 |
| liver | 947 | 1090 | 211 |
| ComBat | 4009 | 180 | 0 |

To further validate the genes selected by other methods, we converted probe IDs to gene symbols and uploaded them to Enrichr. Table 5 presents the results for the other methods on the “Mouse Gene Atlas” category in Enrichr. For muscle, neither SAM nor *t* test could select muscle as top ranked tissues whereas limma could identify only two probes as muscle-specific genes (see Table 4). Thus, the other methods are not better than the proposed method that could identify muscle specificity correctly (Table 3). Figure 2 shows the Venn diagrams between selected genes. Since the proposed method selects different genes from those specifically selected in individual tissues, an integrated analysis is a valuable method.

**Table 5.** Top three terms by other methods in the “Mouse Gene Atlas” category in Enrichr

| Term | Overlap | P-value | Adjusted P-value |
| --- | --- | --- | --- |
| *t* test |  |  |  |
| Adipose |  |  |  |
| adipose brown | 38/456 | $4.17 \times 10^{-12}$ | $3.92 \times 10^{-10}$ |
| mammary gland lact | 12/104 | $3.87 \times 10^{-6}$ | $1.82 \times 10^{-4}$ |
| macrophage peri LPS thio 0 hrs | 18/353 | $1.14 \times 10^{-3}$ | $3.59 \times 10^{-2}$ |
| Muscle |  |  |  |
| adipose brown | 29/456 | $4.34 \times 10^{-26}$ | $2.26 \times 10^{-24}$ |
| heart | 21/568 | $3.88 \times 10^{-14}$ | $1.01 \times 10^{-12}$ |
| mammary gland lact | 4/104 | $1.15 \times 10^{-3}$ | $1.99 \times 10^{-2}$ |
| Liver |  |  |  |
| liver | 90/928 | $2.53 \times 10^{-16}$ | $2.38 \times 10^{-14}$ |
| adipose brown | 40/456 | $9.53 \times 10^{-7}$ | $4.48 \times 10^{-5}$ |
| kidney | 40/554 | $9.11 \times 10^{-5}$ | $2.86 \times 10^{-3}$ |
| ComBat |  |  |  |
| bone marrow | 107/413 | $1.04 \times 10^{-10}$ | $9.98 \times 10^{-9}$ |
| osteoblast day 21 | 75/264 | $8.31 \times 10^{-10}$ | $3.99 \times 10^{-8}$ |
| embryonic stem line V26 2 p16 | 149/728 | $9.44 \times 10^{-7}$ | $3.02 \times 10^{-5}$ |
| sam |  |  |  |
| Adipose |  |  |  |
| adipose brown | 51/456 | $2.61 \times 10^{-16}$ | $2.48 \times 10^{-14}$ |
| mammary gland lact | 12/104 | $5.67 \times 10^{-5}$ | $2.69 \times 10^{-3}$ |
| macrophage peri LPS thio 0 hrs | 23/353 | $3.54 \times 10^{-4}$ | $1.12 \times 10^{-2}$ |
| Muscle |  |  |  |
| adipose brown | 33/456 | $2.16 \times 10^{-29}$ | $1.21 \times 10^{-27}$ |
| heart | 23/568 | $7.01 \times 10^{-15}$ | $1.96 \times 10^{-13}$ |
| mammary gland lact | 4/104 | $1.91 \times 10^{-3}$ | $3.47 \times 10^{-2}$ |
| Liver |  |  |  |
| liver | 93/928 | $3.10 \times 10^{-14}$ | $2.91 \times 10^{-12}$ |
| adipose brown | 43/456 | $1.63 \times 10^{-6}$ | $7.66 \times 10^{-5}$ |
| kidney | 43/554 | $1.76 \times 10^{-4}$ | $5.51 \times 10^{-3}$ |
| Cell cycle | 11/124 | $2.95 \times 10^{-9}$ | $4.71 \times 10^{-7}$ |
| Oocyte meiosis | 9/129 | $6.65 \times 10^{-7}$ | $5.32 \times 10^{-5}$ |
| Progesterone-mediated oocyte maturation | 8/100 | $1.00 \times 10^{-6}$ | $5.34 \times 10^{-5}$ |
| limma |  |  |  |
| Adipose |  |  |  |
| adipose brown | 14/456 | $4.19 \times 10^{-8}$ | $2.85 \times 10^{-6}$ |
| adipose white | 4/199 | $1.61 \times 10^{-2}$ | $5.46 \times 10^{-1}$ |
| intestine small | 6/466 | $2.59 \times 10^{-2}$ | $5.87 \times 10^{-1}$ |
| Liver |  |  |  |
| liver | 33/928 | $2.39 \times 10^{-11}$ | $1.60 \times 10^{-9}$ |
| adipose brown | 7/456 | $1.31 \times 10^{-1}$ | $1.00 \times 10^0$ |
| heart | 8/568 | $1.57 \times 10^{-1}$ | $1.00 \times 10^0$ |

**Figure 2.**
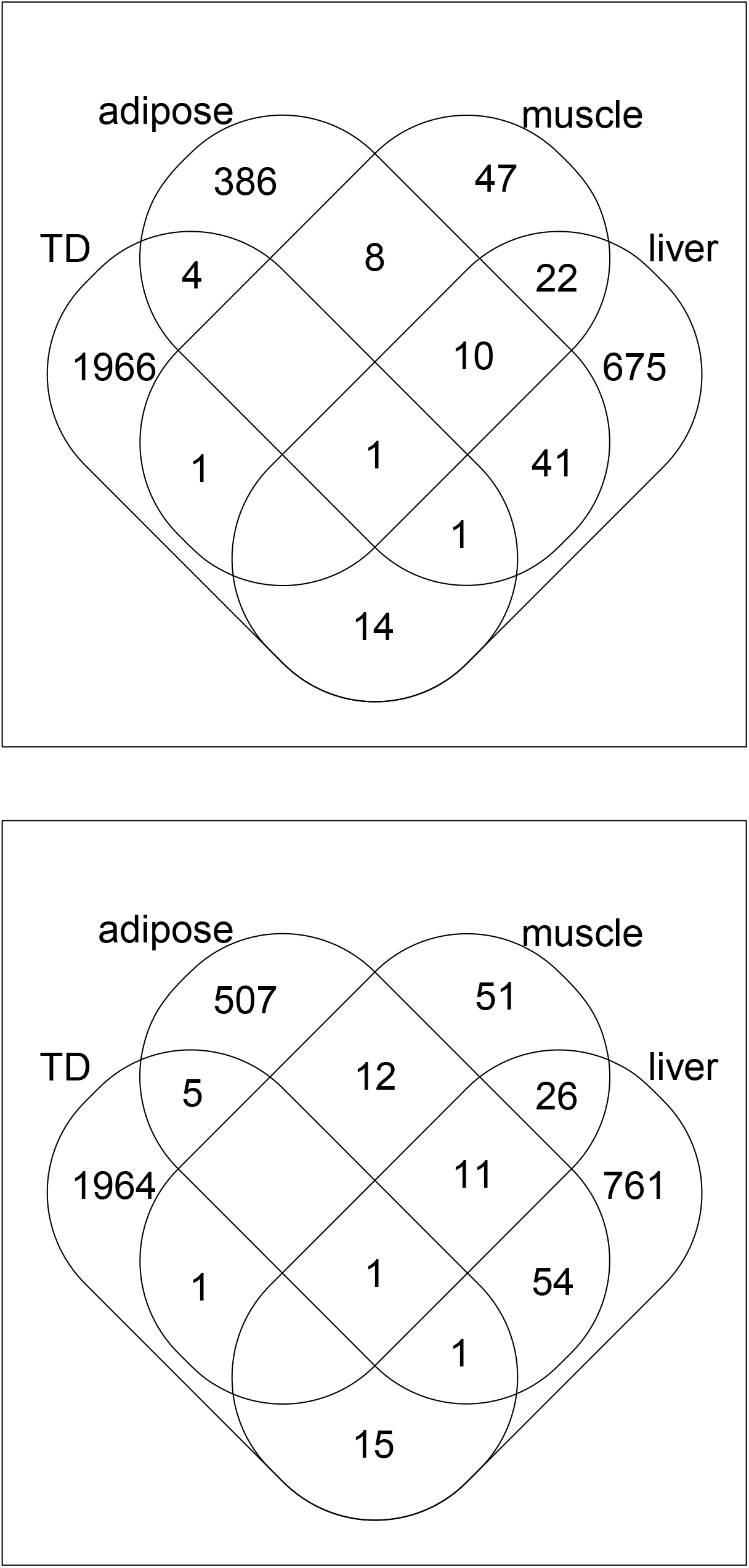
Venn diagrams between genes selected by various methods. Upper: *t* test, lower: SAM.

Finally, based on the genes associated with probes shown in Table 4, we found that the “KEGG 2021 Human” category in Enrichr does not include neurodegenerative diseases (see the supplementary materials). Thus, the association between neurodegenerative diseases and diabetes mellitus can be found only when an integrated analysis, such as the proposed method, is employed. In this sense, an integrated analysis is more than a simple union of individual analysis and can identify factors that cannot be identified by individual analyses, such as potentially associated diseases. Thus, an integrated analysis of gene expression profiles in individual tissues provide more in-depth information than individual analyses, at least for certain cases. Thus, integrated analyses of gene expression profiles in individual tissues should be encouraged.

It may be plausible for other integrated methods to perform similarly. If this is true, the advanced methods that we have proposed here are not required. To rule out this possibility, we apply ComBat [17] to remove the batch effect between the three tissue typed, since we selected genes whose expression is independent of tissues as can be seen in Fig. **??**; Table 4 shows the results. It is seldom reported to be successful. Limma failed to select any DEGs, and the number of genes selected by the *t* test and SAM is markedly different from each other in contrast to the identification of tissue-specific DEGs, whose numbers are more coincident across the three methods (Table 4).

Biological validation is also worse; Table 5 shows the result of the “Mouse Gene Atlas”. None of tissues used in the experiments are listed whereas the proposed method is (Table 3). In addition to this, based on the genes associated with probes shown in Table 4, we found that the “KEGG 2021 Human” category in Enrichr does not include neurodegenerative diseases (see the supplementary materials) that were detected using the proposed method (Table 1). In conclusion, integrated analysis using ComBat is inferior to the proposed method.

One might wonder why an integrated analysis of three tissues from patients with diabetes mellitus can identify associations with neurodegenerative diseases. The PCA and TD-based unsupervised FE methods are frequently able to detect disease associations. We previously identified an association between cancer and amyotrophic lateral sclerosis [18] without investigating cancer gene expression and an association between heart diseases and posttraumatic stress disorder [19] without investigating brain gene expression. Therefore, we were not surprised that the integrated analysis using the proposed method was able to identify disease associations. To our knowledge, few studies have attempted to predict the association between diseases using gene expression, although many studies have focused on the associations between genes and disease [20–22] and between drugs and disease association [23–25]. Our proposed strategy would be useful for such studies.

## 5. Conclusions

In this study, we applied the proposed TD-based unsupervised FE with SD optimization method to perform an integrated analysis of gene expression measured in three distinct tissues using microarray architecture; moreover, the proposed method has not been applied to such data in previous studies. The results show that the proposed method can identify more genes than individual analyses. The selected genes are known to be expressed in all three tissues, and they are also enriched in many neurodegenerative diseases that have a known association with diabetes mellitus but cannot be identified by individual analysis. In this sense, integrated analyses might have the ability to identify additional factors relative to individual analyses.

## Author Contributions

Y.-h.T. planned the research and performed the analyses. Y.-h.T. and T.T. evaluated the results, discussions, and outcomes and wrote and reviewed the manuscript. All authors have read and agreed to the published version of the manuscript.

## Institutional Review Board Statement

Not applicable

## Informed Consent Statement

Not applicable

## Data Availability Statement

All of the data used in this study are available in GEO ID GSE13268, GSE13269, and GSE13270.

## References

1. Taguchi, Y.H. Unsupervised Feature Extraction Applied to Bioinformatics; Springer International Publishing, 2020. doi:10.1007/978-3-030-22456-1.

2. Taguchi, Y.h.; Turki, T. Tensor decomposition- and principal component analysis-based unsupervised feature extraction to select more reasonable differentially expressed genes: Optimization of standard deviation versus state-of-art methods. bioRxiv 2022, [https://www.biorxiv.org/content/early/2022/02/22/2022.02.18.481115.full.pdf]. doi:10.1101/2022.02.18.481115.

3. Taguchi, Y.H.; Turki, T. Principal component analysis- and tensor decomposition-based unsupervised feature extraction to select more reasonable differentially methylated cytosines: Optimization of standard deviation versus state-of-the-art methods. bioRxiv 2022, [https://www.biorxiv.org/content/early/2022/04/05/2022.04.02.486807.full.pdf]. doi:10.1101/2022.04.02.486807.

4. Roy, S.S.; Taguchi, Y.h. Tensor decomposition and principal component analysis-based unsupervised feature extraction outperforms state-of-the-art methods when applied to histone modification profiles. bioRxiv 2022, [https://www.biorxiv.org/content/early doi:10.1101/2022.04.29.490081.

5. Xue, B.; Nie, J.; Wang, X.; DuBois, D.C.; Jusko, W.J.; Almon, R.R. Effects of High Fat Feeding on Adipose Tissue Gene Expression in Diabetic Goto-Kakizaki Rats. Gene Regulation and Systems Biology 2015, 9, GRSB.S25172, [https://doi.org/10.4137/GRSB.S25172]. PMID: 26309393, doi:10.4137/GRSB.S25172.

6. Xie, Z.; Bailey, A.; Kuleshov, M.V.; Clarke, D.J.B.; Evangelista, J.E.; Jenkins, S.L.; Lachmann, A.; Wojciechowicz, M.L.; Kropiwnicki, E.; Jagodnik, K.M.; et al. Gene Set Knowledge Discovery with Enrichr. Current Protocols 2021, 1, e90. [https://currentprotocols.onlinelibrary.wiley.com/doi/pdf/10.1002/cpz1.90]. doi:https://doi.org/10.1002/cpz1.90.

7. Umegaki, H., Neurodegeneration in Diabetes Mellitus. In Neurodegenerative Diseases; Ahmad, S.I., Ed.; Springer US:New York, NY, 2012; pp. 258–265. doi:10.1007/978-1-4614-0653-2_19.

8. Ristow, M. Neurodegenerative disorders associated with diabetes mellitus. Journal of Molecular Medicine 2004, 82. doi:10.1007/s00109-004-0552-1.

9. Nasrolahi, A.; Mahmoudi, J.; Noori-Zadeh, A.; Haghani, K.; Bakhtiyari, S.; Darabi, S. Shared Pathological Mechanisms Between Diabetes Mellitus and Neurodegenerative Diseases. Current Pharmacology Reports 2019, 5, 219–231. doi:10.1007/s40495-019-00191-8.

10. Madhusudhanan, J.; Suresh, G.; Devanathan, V. Neurodegeneration in type 2 diabetes: Alzheimer’s as a case study. Brain and Behavior 2020, 10, e01577. [https://onlinelibrary.wiley.com/doi/pdf/10.1002/brb3.1577]. doi:https://doi.org/10.1002/brb3.1577.

11. León, K.I.L.D.; Bertadillo-Jilote, A.D.; García-Gutiérrez, D.G.; Meraz-Ríos, M.A. Alzheimer’s Disease and Type 2 Diabetes Mellitus: Molecular Mechanisms and Similarities. In Neurodegenerative Diseases; Tunali, N.E., Ed.; IntechOpen: Rijeka, 2020; chapter 4. doi:10.5772/intechopen.92581.

12. Sun, H.; Wang, Y. A new branch connecting thermogenesis and diabetes. Nature Metabolism 2019, 1, 845–846. doi:10.1038/s42255-019-0112-1.

13. Lewis, M.T.; Kasper, J.D.; Bazil, J.N.; Frisbee, J.C.; Wiseman, R.W. Quantification of Mitochondrial Oxidative Phosphorylation in Metabolic Disease: Application to Type 2 Diabetes. International Journal of Molecular Sciences 2019, 20. doi:10.3390/ijms20215271.

14. Holm, L.J.; Mnsted, M.O.; Haupt-Jorgensen, M.; Buschard, K. PPARs and the Development of Type 1 Diabetes. PPAR Res 2020, 2020, 6198628.

15. Tusher, V.G.; Tibshirani, R.; Chu, G. Significance analysis of microarrays applied to the ionizing radiation response. Proceedings of the National Academy of Sciences 2001, 98, 5116–5121, [https://www.pnas.org/content/98/9/5116.full.pdf]. doi:10.1073/pnas.091062498.

16. Ritchie, M.E.; Phipson, B.; Wu, D.; Hu, Y.; Law, C.W.; Shi, W.; Smyth, G.K. limma powers differential expression analyses for RNA-sequencing and microarray studies. Nucleic Acids Research 2015, 43, e47.#x2013;e47, [https://academic.oup.com/nar/article-pdf/43/7/e47/7207289/gkv007.pdf]. doi:10.1093/nar/gkv007.

17. Johnson, W.E.; Li, C.; Rabinovic, A. Adjusting batch effects in microarray expression data using empirical Bayes methods. Biostatistics 2006, 8, 118–127, [https://academic.oup.com/biostatistics/article-pdf/8/1/118/25435561/kxj037.pdf]. doi:10.1093/biostatistics/kxj037.

18. Taguchi, Y.H.; Wang, H. Genetic Association between Amyotrophic Lateral Sclerosis and Cancer. Genes 2017, 8. doi:10.3390/genes8100243.

19. Taguchi, Y.H.; Iwadate, M.; Umeyama, H. Principal component analysis-based unsupervised feature extraction applied to in silico drug discovery for posttraumatic stress disorder-mediated heart disease. BMC Bioinformatics 2015, 16. doi:10.1186/s12859-015-0574-4.

20. Babbi, G.; Martelli, P.L.; Profiti, G.; Bovo, S.; Savojardo, C.; Casadio, R. eDGAR: a database of Disease-Gene Associations with annotated Relationships among genes. BMC Genomics 2017, 18. doi:10.1186/s12864-017-3911-3.

21. Luo, P.; Xiao, Q.; Wei, P.J.; Liao, B.; Wu, F.X. Identifying Disease-Gene Associations With Graph-Regularized Manifold Learning. Frontiers in Genetics 2019, 10. doi:10.3389/fgene.2019.00270.

22. Opap, K.; Mulder, N. Recent advances in predicting gene?disease associations [version 1; peer review: 2 approved]. F1000Research 2017, 6. doi:10.12688/f1000research.10788.1.

23. Huang, F.; Qiu, Y.; Li, Q.; Liu, S.; Ni, F. Predicting Drug-Disease Associations via Multi-Task Learning Based on Collective Matrix Factorization. Frontiers in Bioengineering and Biotechnology 2020, 8. doi:10.3389/fbioe.2020.00218.

24. Jiang, H.; Huang, Y. An effective drug-disease associations prediction model based on graphic representation learning over multi-biomolecular network. BMC Bioinformatics 2022, 23. doi:10.1186/s12859-021-04553-2.

25. Yu, Z.; Huang, F.; Zhao, X.; Xiao, W.; Zhang, W. Predicting drug–disease associations through layer attention graph convolutional network. Briefings in Bioinformatics 2020, 22, [https://academic.oup.com/bib/article-pdf/22/4/bbaa243/39135298/bbaa243.pdf]. bbaa243, doi:10.1093/bib/bbaa243.

