## Supplementary material for "Integrated Analysis of Tissue-specific Gene Expression in Diabetes by Tensor Decomposition Can Identify Possible Associated Diseases": Sample.R

```
#Currwnt directory should includ the following files
#GSE13268_series_matrix.txt.gz
#GSE13269_series_matrix.txt.gz
#GSE13270_series_matrix.txt.gz

require(rTensor)
x <- read.csv("GSE13268_series_matrix.txt.gz",sep="\t",comment.char="!")
x1 <- read.csv("GSE13269_series_matrix.txt.gz",sep="\t",comment.char="!")
x2 <- read.csv("GSE13270_series_matrix.txt.gz",sep="\t",comment.char="!")
Z <- array(NA,c(dim(x)[1],5,5,2,2,3))
for (i in c(1:5)){
    cat(i, " ")
    Z[,i,1:5,1,1,1]<-data.matrix(x[,-1][,20*(i-1)+c(1:5)])
    Z[,i,1:5,2,1,1]<-data.matrix(x[,-1][,20*(i-1)+c(6:10)])
    Z[,i,1:5,1,2,1]<-data.matrix(x[,-1][,20*(i-1)+c(11:15)])
    Z[,i,1:5,2,2,1]<-data.matrix(x[,-1][,20*(i-1)+c(16:20)])
}
for (i in c(1:5)){
    cat(i, " ")
    Z[,i,1:5,1,1,2]<-data.matrix(x1[,-1][,20*(i-1)+c(1:5)])
    Z[,i,1:5,2,1,2]<-data.matrix(x1[,-1][,20*(i-1)+c(6:10)])
    Z[,i,1:5,1,2,2]<-data.matrix(x1[,-1][,20*(i-1)+c(11:15)])
    Z[,i,1:5,2,2,2]<-data.matrix(x1[,-1][,20*(i-1)+c(16:20)])
}
for (i in c(1:5)){
    cat(i, " ")
    Z[,i,1:5,1,1,3]<-data.matrix(x2[,-1][,20*(i-1)+c(1:5)])
    Z[,i,1:5,2,1,3]<-data.matrix(x2[,-1][,20*(i-1)+c(6:10)])
    Z[,i,1:5,1,2,3]<-data.matrix(x2[,-1][,20*(i-1)+c(11:15)])
    Z[,i,1:5,2,2,3]<-data.matrix(x2[,-1][,20*(i-1)+c(16:20)])
}
# gene,week, repl, treated vs control, Goto vs W

Z <- apply(Z,2:6,scale)
HOSVD <- hosvd(as.tensor(Z),c(10,5,5,2,2,3))
HOSVD$Z@data[1:10,2,1,2,1,1]
#[1]   3.505367   8.092320 -15.707937   9.134575   6.723113   6.227675   6.890919
#[8]  -3.287872  -1.529140  -9.158979

th <- function(sd){
    P2 <- pchisq((HOSVD$U[[1]][,3]/sd)^2,1,lower.tail=F)
    hc<- hist(1-P2,breaks=100,plot=F)
    return(sd(hc$count[1:sum(hc$breaks<1-min(P2[p.adjust(P2,"BH")>0.01]))]))
}
sd <- optim(0.01,th)$par
P <- pchisq((HOSVD$U[[1]][,3]/sd)^2,1,lower.tail=F)
print(c(sd,sd(HOSVD$U[[1]][p.adjust(P,"BH")>0.01,3])))
#[1] 0.001514603 0.001206368

pdf(file="hist.pdf")
hist(1-P,breaks=100,cex.axis=2,cex.lab=1.5)
dev.off()

table(p.adjust(P,"BH")<0.01)
#FALSE  TRUE 
#28557  2542 

gene <- x[p.adjust(P,"BH")<0.01,1]
write.table(file="gene.csv",gene,sep="\t",col.names=F,row.names=F,quote=F)

jj<-1 #adipose
#jj<-2 #muscle
#jj<-3 #liver 
X <- Z[,,,,,jj]
X1 <- X[,,,1,]
dim(X1) <- c(31099,50)
X2 <- X[,,,2,]
dim(X2) <- c(31099,50)
XX <- cbind(X1,X2)

#sam
require(siggenes)
SAM <- sam(XX,rep(0:1,each=50))
table(SAM@q.value<0.01)

#jj<-1 adipose
#FALSE  TRUE 
#30326   773 

#jj<-2 muscle
#FALSE  TRUE 
#30980   119

#jj<-3 liver
#FALSE  TRUE 
#30009  1090 

gene <- x[SAM@q.value<0.01,1]
write.table(file="gene_sam_adipose.csv",gene,sep="\t",col.names=F,row.names=F,quote=F)
write.table(file="gene_sam_muscle.csv",gene,sep="\t",col.names=F,row.names=F,quote=F)
write.table(file="gene_sam_liver.csv",gene,sep="\t",col.names=F,row.names=F,quote=F)

#limma
require(limma)
time <- array(NA,dim(Z[,,,,,jj])[-1])
for (i in c(1:5)) {time[i,,,]<-i}
time1 <- time[,,1,]
dim(time1) <- 50
time2 <- time[,,2,]
dim(time2) <- 50
time <- c(time1,time2)
targets <- data.frame(Group=factor(rep(1:2,each=50)),Time=time)
library(splines)
T <- ns(targets$Time, df=3)
Group <- factor(targets$Group)
design <- model.matrix(~Group*T)
fit <- lmFit(XX, design)
fit <- eBayes(fit)
TABLE <- topTable(fit, coef=6:8,numbe=dim(XX)[1])
table(TABLE$adj.P.Val<0.01)
#jj<-1 adipose
#FALSE  TRUE 
#30983   116 

#jj<-2 muscle
#FALSE  TRUE 
#31097     2 


#jj<-3 liver
#FALSE  TRUE 
#30888   211 


gene <- x[rownames(TABLE)[TABLE$adj.P.Val<0.01],1]
write.table(file="gene_limma_adipose.csv",gene,sep="\t",col.names=F,row.names=F,quote=F)
write.table(file="gene_limma_liver.csv",gene,sep="\t",col.names=F,row.names=F,quote=F)

#t test
P <- apply(XX,1,function(x){t.test(x~rep(1:2,each=50))$p.value})
table(p.adjust(P,"BH")<0.01)

#jj<-1 adipose
#FALSE  TRUE 
#30533   566 

#jj<-2 muscle
#FALSE  TRUE 
#30999   100 

#jj<-3 liver
#FALSE  TRUE 
#30152   947 

gene <- x[p.adjust(P,"BH")<0.01,1]
write.table(file="gene_ttest_adipose.csv",gene,sep="\t",col.names=F,row.names=F,quote=F)
write.table(file="gene_ttest_muscle.csv",gene,sep="\t",col.names=F,row.names=F,quote=F)
write.table(file="gene_ttest_liver.csv",gene,sep="\t",col.names=F,row.names=F,quote=F)


#combat

require(neuroCombat)
data.harmonized <- neuroCombat(dat=cbind(x[,-1],x1[,-1],x2[,-1]), batch=rep(1:3,each=101))
#limma
require(limma)
targets3 <- rbind(targets,targets,targets)
library(splines)
T <- ns(targets3$Time, df=3)
Group <- factor(targets3$Group)
design <- model.matrix(~Group*T)
XX <- data.harmonized$dat.standardized[,-c(101,202,303)]
fit <- lmFit(XX, design)
fit <- eBayes(fit)
TABLE <- topTable(fit, coef=6:8,numbe=dim(XX)[1])
table(TABLE$adj.P.Val<0.01)
#FALSE 
#31099 

#SAM
SAM <- sam(XX,rep(rep(0:1,each=50),3))
table(SAM@q.value<0.01)
#FALSE  TRUE 
#30919   180

#t test
P <- apply(XX,1,function(x){t.test(x~rep(rep(1:2,each=50),3))$p.value})
table(p.adjust(P,"BH")<0.01)
#FALSE  TRUE 
#27090  4009 
```
