## Supplementary material for "Integrated Analysis of Tissue-specific Gene Expression in Diabetes by Tensor Decomposition Can Identify Possible Associated Diseases": Supplementary_Information.pdf

#### 1 The brief description of the HOSVD algorithm

For demonstration, consider the tensor $x_{ijk} \in \mathbb{R}^{N\times M\times K}$. HOSVD is an algorithm that applies SVD to the unfolded tensor, either $x_{i(jk)} \in \mathbb{R}^{N\times MK}$, $x_{j(ik)} \in \mathbb{R}^{M\times NK}$, or $x_{k(ij)} \in \mathbb{R}^{K\times NM}$,

$$x_{i(jk)} = \sum_{\ell_1=1}^{\min(N,MK)} \lambda_{\ell_1} u_{\ell_1 i} v_{\ell_1 jk} \quad (1)$$

$$x_{j(ik)} = \sum_{\ell_2=1}^{\min(M,NK)} \lambda_{\ell_2} u_{\ell_2 j} v_{\ell_2 ik} \quad (2)$$

$$x_{k(ij)} = \sum_{\ell_3=1}^{\min(K,NM)} \lambda_{\ell_3} u_{\ell_3 k} v_{\ell_3 ij} \quad (3)$$

$$\quad (4)$$

where $u_{\ell_1 i} \in \mathbb{R}^{\min(N,MK)\times N}$, $v_{\ell_1 jk} \in \mathbb{R}^{\min(N,MK)\times MK}$, $u_{\ell_2 j} \in \mathbb{R}^{\min(M,NK)\times M}$, $v_{\ell_2 ik} \in \mathbb{R}^{\min(M,NK)\times NK}$, $u_{\ell_3 k} \in \mathbb{R}^{\min(K,MM)\times K}$, $v_{\ell_3 ij} \in \mathbb{R}^{\min(K,NM)\times NM}$. A core tensor $G(\ell_1\ell_2\ell_3)$ can be calculated as

$$G(\ell_1\ell_2\ell_3) = \sum_{i=1}^{N}\sum_{j=1}^{M}\sum_{k=1}^{K} x_{ijk} u_{\ell_1 i} u_{\ell_2 j} u_{\ell_3 k}. \quad (5)$$

Subsequently, we obtain the following

$$x_{ijk} = \sum_{\ell_1=}^{N}\sum_{\ell_2=1}^{M}\sum_{\ell_3=1}^{K} G(\ell_1\ell_2\ell_3) u_{\ell_1 i} u_{\ell_2 j} u_{\ell_3 k} \quad (6)$$

where $u_{\ell_1 i} \in \mathbb{R}^{N\times N}$, $u_{\ell_2 j} \in \mathbb{R}^{M\times M}$, and $u_{\ell_3 k} \in \mathbb{R}^{K\times K}$. When $u_{\ell_1 i}$, $u_{\ell_2 j}$, and $u_{\ell_3 k}$, are smaller than those computed in eqs. (1), (2), and (3), missing values are filled with zero. Then, $u_{\ell_1 i}$, $u_{\ell_2 j}$, and $u_{\ell_3 k}$ are the resultant orthogonal matrices obtained.

#### 2 TD-based unsupervised FE with SD optimization

HOSVD was applied to $x_{ijkmst}$, and we obtained the following:

$$x_{ijkmst} = \sum_{\ell_1=1}^{5}\sum_{\ell_2=1}^{5}\sum_{\ell_3=1}^{2}\sum_{\ell_4=1}^{2}\sum_{\ell_5=1}^{3}\sum_{\ell_6=1}^{31099} G(\ell_1\ell_2\ell_3\ell_4\ell_5\ell_6) u_{\ell_1 j} u_{\ell_2 k} u_{\ell_3 m} u_{\ell_4 s} u_{\ell_5 t} u_{\ell_6 i} \tag{7}$$

where $G(\ell_1\ell_2\ell_3\ell_4\ell_5\ell_6) \in \mathbb{R}^{5\times5\times2\times2\times3\times31099}$ is a core tensor and $u_{\ell_1 j} \in \mathbb{R}^{5\times5}$, $u_{\ell_2 k} \in \mathbb{R}^{5\times5}$, $u_{\ell_3 m} \in \mathbb{R}^{2\times2}$, $u_{\ell_4 s} \in \mathbb{R}^{2\times2}$, $u_{\ell_5 t} \in \mathbb{R}^{3\times3}$, and $u_{\ell_6 i} \in \mathbb{R}^{31099\times31099}$ are singular value matrices and orthogonal matrices.

After identifying $u_{\ell_1 j}$, $u_{\ell_2 k}$, $u_{\ell_3 m}$, $u_{\ell_4 s}$, and $u_{\ell_5 t}$ of interest, we attempted to identify $G(\ell_1\ell_2\ell_3\ell_4\ell_5\ell_6)$ with the largest absolute value and fixed $\ell_1$ to $\ell_5$. Then, we attributed $P$-values to $i$ by assuming that $u_{\ell_6 i}$ obeys a Gaussian distribution as follows:

$$P_i = P_{\chi^2}\left[ > \left(\frac{u_{\ell_6 i}}{\sigma_{\ell_6}}\right)^2 \right] \tag{8}$$

where $P_{\chi^2}[> x]$ is the cumulative $\chi^2$ distribution under the assumption that it is larger than $x$ and $\sigma_{\ell_6}$ is the SD. $P_i$ is corrected with the BH criterion [?].

The SD is optimized so that $u_{\ell_6 i}$ obeys the Gaussian distribution (null hypothesis) as much as possible [?], and the optimization process is as follows. Initially, we compute the histogram $h_p$, which includes the number of $i$ values that satisfy the following:

$$\frac{p}{N_p} \le 1 - \text{adjusted}P_i \le \frac{p+1}{N_p}, 0 \le p \le N_p - 1 \tag{9}$$

Then, the standard deviation of $h_p$ for adjusted$P_i > p_0$ for the threshold value $p_0$,

$$\Delta h_p = \frac{1}{N'_p} \sum_{\text{adjusted}\frac{p}{N_P} > p_0} (h_p - \langle h_p \rangle)^2 \tag{10}$$

$$\langle h_p \rangle = \frac{1}{N'_p} \sum_{\text{adjusted}\frac{p}{N_P} > p_0} h_p \tag{11}$$

$$N'_P = \sum_{\text{adjusted}\frac{p}{N_P} > p_0} 1 \tag{12}$$

is minimized. Therefore, for $1 - \text{adjusted}P_i < 1 - p_0$, $h_p$ takes a constant value (i.e., it obeys the null hypothesis) as much as possible, whereas for $1 - \text{adjusted}P_i > 1 - p_0$, $h_p$ presents a peak value.

The $P_i$ values are finally calculated using the optimized SD, and $i$ values associated with adjusted $P$-values less than 0.01 are selected. Probes, $i$s, are converted to gene symbols using the gene ID conversion tool in DAVID [?].

The genes selected are presumed to be coincident with the profiles associated with selected $u_{\ell_1 j}$, $u_{\ell_2 k}$, $u_{\ell_3 m}$, $u_{\ell_4 s}$, and $u_{\ell_5 t}$; this corresponds to the identification of differentially expressed genes.

For details, check sample R source code in supplementary materials.

### 3 Gene sepections

The results indicated that $u_{2j}$ represents the dependence on time, $u_{2m}$ represents the distinction between the controls and treatment, $u_{1k}$ represents the independence of the replicate, $u_{1s}$ represents the independence of strains, and $u_{1t}$ represents the independence of tissues (Fig. 1). Table 1 presents

Table 1: The core tensor defined in eq. (7) when HOSVD is applied to data sets, $G(2,1,2,1,1,\ell_6)$.

| $\ell_6$ | $G(2,1,2,1,1,\ell_6)$ | $\ell_6$ | $G(2,1,2,1,1,\ell_6)$ |
|---|---|---|---|
| 1 | 3.505367 | 6 | 6.227675 |
| 2 | 8.092320 | 7 | 6.890919 |
| 3 | -15.707937 | 8 | -3.287872 |
| 4 | 9.134575 | 9 | -1.529140 |
| 5 | 6.723113 | 10 | -9.158979 |

$G(2,1,2,1,1,\ell_6)$, and it shows that $G(2,1,2,1,1,3)$ has the highest absolute value. Then, $u_{3i}$ (i.e., $\ell_6 = 3$) is used to select probes. The $P_i$ values are determined using eq. (8), with $\ell_6 = 3$. SD optimization was performed, and $\sigma_3 = 0.001514603$. A histogram of 1-$P_i$ after SD optimization is presented in Figure 2 and the histogram is more similar to that of the null hypothesis, i.e., the combination of a flat region and a sharp peak. Then, the $P$-values corresponding to the $i$th probe are determined with eq. (8), and the $P_i$ values are corrected by BH criterion. Finally, 2,452 selected probes are associated with adjusted $P$-values less than 0.01, and they are further converted to 2,281 gene symbols.

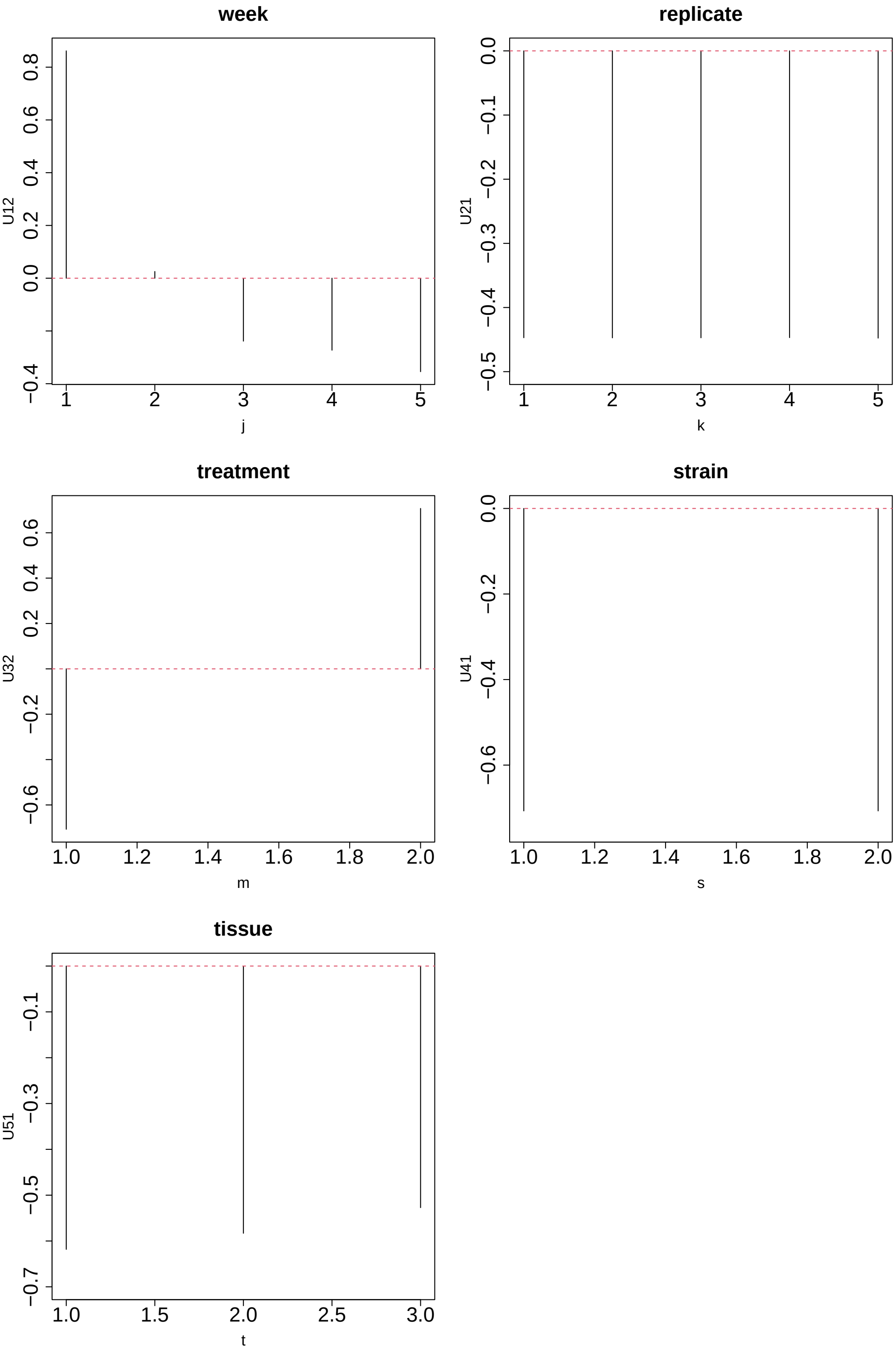


Figure 1: Singular value vectors defined in eq. (7) when HOSVD is applied to data sets. $u_{2j}$ (week), $u_{1k}$ (replicate), $u_{2m}$ (treatment), $u_{1s}$ (strain), and $u_{1t}$ (tissue).

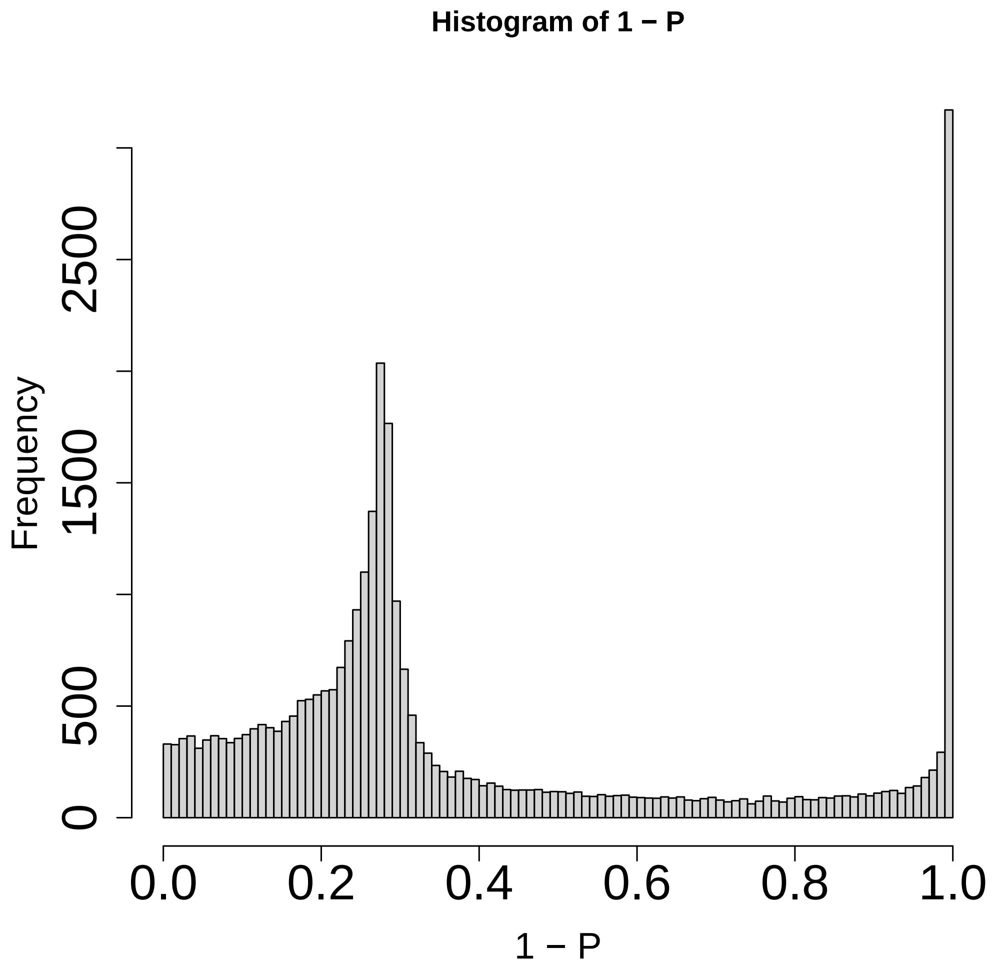


Figure 2: Histogram of 1-$P_i$.
